# MaxTiC: Fast ranking of a phylogenetic tree by Maximum Time Consistency with lateral gene transfers

**DOI:** 10.1101/127548

**Authors:** Cédric Chauve, Akbar Rafiey, Adrián A. Davín, Celine Scornavacca, Philippe Veber, Bastien Boussau, Gergely J. Szöllősi, Vincent Daubin, Eric Tannier

**Affiliations:** Department of Mathematics, Simon Fraser University, Burnaby (BC), Canada; Inria Grenoble Rhône-Alpes, F-38334 Montbonnot, France; Univ Lyon, Université Lyon 1, CNRS, Laboratoire de Biométrie et Biologie Évolutive UMR5558, F-69622 Villeurbanne, France; Institut des Sciences de l’Evolution, Université de Montpellier, CNRS, IRD, EPHE 34095 Montpellier Cedex 5, France; MTA-ELTE “Lendulet” Evolutionary Genomics Research Group, Budapest Hungary; Deptartment of Biological Physics, Eötvos Loránd University, Budapest Hungary

## Abstract

Lateral gene transfers between ancient species contain information about the relative timing of species diversification. Specifically, the ancestors of a donor species must have existed before the descendants of the recipient species. Hence, the detection of a transfer event can be translated into a time constraint between nodes of a phylogeny if the donor and recipient can be identified. When a set of transfers is detected by interpreting the phylogenetic discordance between gene trees and a species tree, the set of all deduced time constraints can be used to rank the species tree, *i.e.* order totally its internal nodes. Unfortunately lateral gene transfer detection is challenging and current methods produce a significant proportion of false positives. As a result, often, no ranking of the species tree is compatible with the full set of time constraints deduced from predicted transfers. Here we propose a method, implemented in a software called MaxTiC (Maximum Time Consistency), which takes as input a species tree and a series of (possibly inconsistent) time constraints between its internal nodes, weighted by confidence scores. MaxTiC outputs a ranked species tree compatible with a subset of constraints with maximum cumulated confidence score. We extensively test the method on simulated datasets, under a wide range of conditions that we compare to measures on biological datasets. In most conditions the obtained ranked tree is very close to the real one, confirming the potential of dating the history of life with transfers by maximizing time consistency. MaxTiC is freely available, distributed along with a documentation and several examples: https://github.com/ssolo/ALE/tree/master/maxtic.

## I. Introduction

*Telling the evolutionary time* is usually achieved by combining molecular clocks and the fossil record (Donoghue and Smith, 2003). It was pointed out by Gogarten et al. (1999) and demonstrated by Szöllősi et al. (2012) that there exists a third source of information on evolutionary time in ancient lateral gene transfers.

Indeed, suppose that ancient species *A* transferred a gene to another species *B*, and the latter has descendants that are sampled in a phylogenetic study. If we call *X* the most recent node of this phylogeny that is an ancestor of *A*, and *Y* the node that directly descends from *B*, then *X* must be older than *Y* since a gene from a descendant of *X* has been transferred to an ancestor of *Y* (see Figure 1).

**Figure 1:**
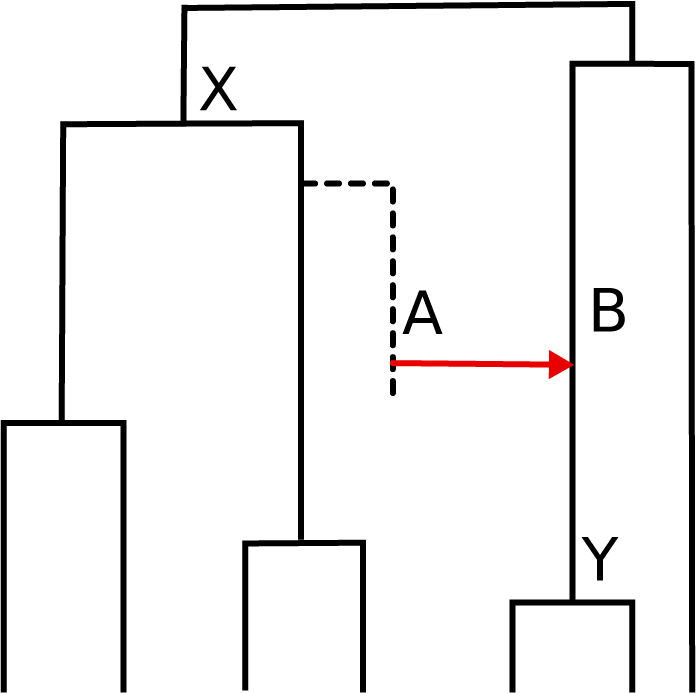
From lateral gene transfer to time constraint. A species tree is depicted, with a transfer from species A to the contemporaneous species B. The donor species A possibly belongs to a lineage with no sampled descendants (dotted line in the phylogeny) (Szöllősi et al., 2013b). The transfer from A to B implies that speciation X must be older than speciation Y. This precedence relation between X and Y constitutes the time constraint associated with the transfer from A to B.

A single transfer can thus provide a time constraint between two nodes of a phylogeny, so many transfers combined can provide a multitude of time constraints that can be used to determine the time order of the internal nodes of a phylogeny and obtain a *ranked phylogeny* (Semple and Steel, 2003). Ranked trees provide a relative timing of diversification. They can be used for example to detect events of diversification burst and to relate them with known events in the history of life (Ford et al., 2009). As lateral gene transfers have probably been very frequent in evolution, in particular among microbes (Ochman et al., 2000), this could constitute the most abundant source of information for dating in the history of life. Interestingly, it may be most plentiful in taxa where fossils are absent.

While dating with transfers is possible in theory, it has rarely been put in pratice, mainly because of the difficulty of detecting lateral gene transfers and identifying the donor and recipient lineages (Ravenhall et al., 2015). The only published attempt to date a phylogeny using a collection of predicted transfers for dating a species tree is the method by Szöllősi et al. (2012), which consists in finding the ranked tree that maximizes the likelihood according to a model of gene tree-species tree reconciliation taking lateral gene transfers into account. Due to the size of the space of dated trees and the time complexity of the gene tree-species tree reconciliation, this method it is computationally demanding and hardly scales up to large datasets. This calls for methodological improvements.

A fast alternative is to detect transfers on an unranked species tree, and combine all transfers to output a ranking. Several programs are available to detect transfers using phylogenetic incongruence between species trees and gene trees without the need of a ranked species tree (Bansal et al., 2012; Stolzer et al., 2012; Szöllősi et al., 2015; Badescu et al., 2016; Jacox et al., 2016). However all these methods output, by construction, sets of transfers that can be *inconsistent*, *i.e.* not compatible with any ranking of the species tree. At most some of them can output time consistent sets for a single gene family (Stolzer et al., 2012; Jacox et al., 2016), possibly at the cost of a high computing time. This inconsistency is due to errors in trees or reconciliations: for example, the species tree can be partly incorrect, gene families may be wrongly inferred, the gene trees are prone to the usual reconstruction uncertainties or systematic artifacts, and reconciliation models are lacking important events such as incomplete lineage sorting, gene conversion and transfers with replacement of an homologous gene (Chan et al., 2017; Hasic and Tannier, 2017a,b).

In this paper we propose a method and an associated tool, called MaxTiC, for *Maximal Time Consistency*, to compute a ranking of an input species tree, given a set of time constraints between internal nodes of the species tree, deduced from lateral gene transfers. The constraints may be weighted with a confidence score, as it is the case in the output of several programs which sample in the space of solutions (Szöllősi et al., 2013a; Szöllősi et al., 2015; Jacox et al., 2016). The output ranked tree aims at maximizing the sum of the confidence scores of the compatible transfers, or equivalently minimizing the incompatible ones. We show that our problem is equivalent to a well-known difficult problem in computer science, the Feedback Arc Set, and propose a method combining three heuristics inspired from the computer science literature on this problem. These are implemented in the MaxTiC tool.

We provide a proof of principle that this method is able to efficiently date phylogenetic trees by generating a number of simulated datasets with SimPhy (Mallo et al., 2016) and detecting transfers with ALE_undated (Szöllősi et al., 2015). We use a wide range of transfer rates, population sizes (which affects the amount of gene tree species tree incongruence due to incomplete lineage sorting, which is not modeled by ALE_undated), variations in the species trees and test the limits of our method. We show that under most conditions tested in our simulations, including some settings with features comparable to the ones observed in published fungi and cyanobacteria datasets, the ranked tree recovered by the method is very close to the true one, but is never exactly the same. Still, this is not due to the heuristic optimization but rather to false transfers inferred by ALE_undated. Indeed, inferred solutions are slightly better, according to the cost function, than the true ranking.

The organization of the paper is the following: We first describe the protocol, including simulations, transfer detection and the conversion of each transfer into a time constraint. Then, we describe our main algorithm and its properties. We finally present the results on the simulated datasets and discuss the possibility to date a tree of life using transfers.

## II. Construction of the simulated datasets

### Simulation by SimPhy

We generated simulated datasets with SimPhy (Mallo et al., 2016), a tool that was developed by an independent team with other purposes than to test our method. This has in particular the consequence of simulating processes that are not considered by our inference method, such as transfers with replacements or incomplete lineage sorting. However, SimPhy has been developed to validate evolutionary inference methods in general, it uses birth-death processes just as our inference method, and it assumes that there is no uncertainty in gene family clustering, which is a limit to independence (Biller et al., 2016). For all sets of parameters, we used SimPhy to generate a ranked species tree with 500 leaves. Along this species tree, we generated 100 to 5000 gene trees with a population size between 2 and 10^6^ individuals per species, null rates of duplications and losses, and a rate of transfers from 10^−9^ to 10^−5^. Note that transfers in SimPhy are actually transfers with replacements, which can be interpreted as a transfer and a loss, so a null loss rate in SimPhy is in fact a loss rate equal to the transfer rate if it is seen from the inference model point of view. The parameters range was chosen to provide a broad overview of how the method performs in various contexts: ranging from too few transfers to settings with too many transfers; from perfect gene trees to very noisy gene trees. Also we explored specific questions, such as "what is the robustness to uncertainties in the species tree, or in gene trees?", or "how is the ranking signal correlated to the number of gene families?". We discuss these points in the Results section. Of course not all combinations of parameters, conditions and questions are tested in this paper. For example, sensitivity to errors in the species tree was only tested with one transfer rate—we chose the one giving a number of transfers close to what we measured on biological datasets. Duplications and losses rates are null, which simulates a filter of the families with universal single copy genes.

### Species extinction

We pruned each leaf of the species tree with a probability 0.8, so that the final species tree has approximately 100 leaves. Gene trees were pruned accordingly by discarding leaves belonging to the removed species. This simulates a sampling of sequenced species, accounting for species extinction or species absence in the study (Szöllősi et al., 2013b).

### Detection of transfers

Transfers were detected by ALEml_undated, a program from the ALE suite (Szöllősi et al., 2013a; Szöllősi et al., 2015). ALEml_undated as input an unranked rooted species tree and an unrooted gene tree, and produces a sample of 100 reconciled gene trees (for each of the simulated gene families), sampled according to their likelihood under a model of duplication, loss, and transfers. Duplication, transfer and loss rates are estimated for each gene family independently with a maximum likelihood algorithm, and the 100 reconciled gene trees are sampled according to these ML rates. We ran ALEml_undated with the (undated) species tree and the gene trees (considered as unrooted) generated by SimPhy.

### From transfers to constraints

In order to reduce the noise from transfers found with a low probability, only transfers found in at least 5% of the reconciliations for one gene family were considered. For each remaining transfer, the most recent node in the species phylogeny which is an ancestor of the donor species is called *X*, and the first descendant node of the recipient species is called *Y*. A constraint is inferred as *X* → *Y*, which means *X* should be older than *Y* (see Figure 1). We assign to the constraint *X* → *Y* the support of the transfer, which is the frequency at which the transfer is found in the 100 reconciled gene trees, summed across all gene families.

### Measuring error in time order reconstruction

In order to compare the true (simulated) ranked tree with the obtained ranked tree we compute a similarity measure derived from the Kendall *τ* distance between total orders. The Kendall distance between two orders is the number of pairs *i*, *j* of elements of the two orders such that *i* is before *j* in one order, and *j* is before *i* in the other. We apply the Kendall distance to total orders of internal nodes of ranked species trees. We normalize this number by the maximum possible Kendall distance given that the two orders have to be derived from the same species tree, to get a number between 0 and 1 (0 for the maximum distance and minimum similarity between orders given a species tree, 1 for two equal orders). This maximum Kendall distance is computed thanks to the following property.

#### Property 1

Given a rooted tree T inducing a partial order P on its internal nodes, two depth first searches of T ordering the children of any node in lexicographical order, respectively and anti-lexicographical order, output two linear extensions of P such that their Kendall distance is maximum, among all pairs of linear extensions of P.

This property is easy to demonstrate: take any pair *i*, *j* of internal nodes of a rooted tree. Either one is the ancestor of the other—and they appear in the same order in any pair of linear extensions—or they are incomparable, with a last common ancestor *a*, having children *a*_1_, the ancestor of *i*, and *a*_2_, the ancestor of *j*. In one depth first search *a*_1_ and its descendants, including *i*, appear before *a*_2_ and its descendants, including *j*, and in the other the opposite holds. So all incomparable pairs appear in a different order, contributing to the Kendall distance. This obviously gives the maximum possible Kendall distance. Our Normalized Kendall similarity between two ranked trees is then:

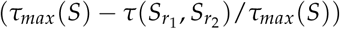

where *τ_max_*(*S*) is the maximum Kendall distance given an unranked species tree *S* and *τ*(*S*_*r*1_, *S*_*r*2_) is the Kendall distance between the two ranked trees.

### Availability

The implementation is available at https://github.com/ssolo/ALE/tree/master/maxtic, along with a documentation, and example datasets.

## III. Finding a maximum consistent set of constraints with MaxTiC

### Definition

We suppose we have as input a rooted, unranked, species tree *S* and a set of weighted constraints *C*, which are directed pairs *X* → *Y* of internal nodes of *S*. We call a constraint *informative* if its two nodes are not related by an ancestor/descendant relationship, and we suppose without loss of generality that *C* contains only informative constraints.

Some constraints might be conflicting, for example like in Figure 2: *Y* is found to be older than *X*, and *Z* is found to be older than *T*, but *T* is an ancestor of *Y* and *X* is an ancestor of *Z*. The two constraints *Y* → *X* and *Z* → *T* cannot be true at the same time in the context of the drawn species tree.

**Figure 2:**
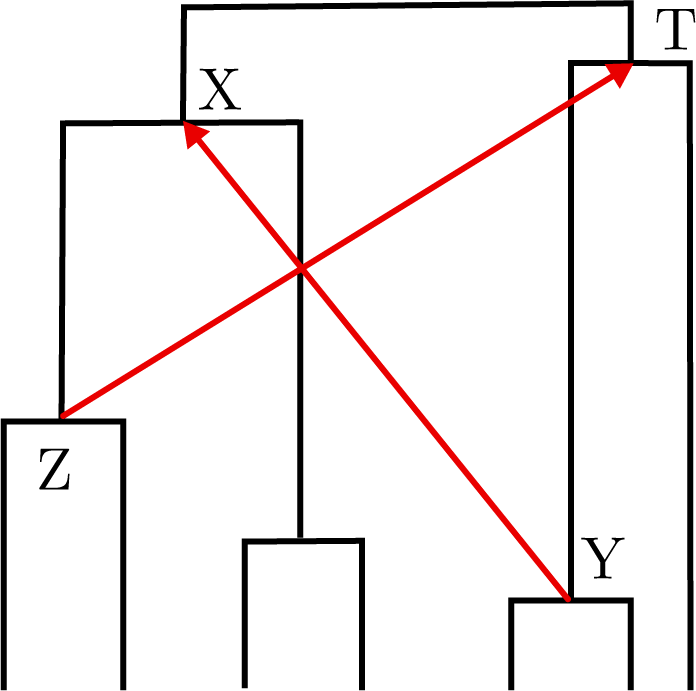
A set of two conflicting constraints. Each of the constraints Y → X and Z → T can be fulfilled by some ranked version of the species tree, but not both.

The problem, that we call *Maximum Time Consistency*, is to find a *ranked* species tree, that is, a total order of the internal nodes compatible with the partial order given by the ancestor/descendant relation in the tree topology. We say that a constraint is *compatible* with a ranked tree if it is directed from an older to a younger node. A subset of constraints on the internal nodes of a tree *S* is *consistent* if there exists a ranked tree based on *S* such that all constraints from this subset are compatible with the ranked tree. Otherwise it is *conflicting*. We search for a maximum weight consistent subset of *C*, which is equivalent to finding a ranked tree with which a maximum weight subset of *C* is compatible.

### Relation with the Feedback Arc Set

If we see the branches of the unranked species tree as arcs of a directed graph with infinite weight, and the constraints as weighted arcs in this graph, then the Maximum Time Consistency problem translates exactly into an instance of the Feedback Arc Set problem. This classical problem is known to be computationally hard: it is NP-complete (Garey and Johnson, 1990), and no approximation with a constant factor is known. Approximation algorithms with *O*(log *n* log log *n*) factors exist (Even et al., 1998), however the best algorithms to solve it in practice have been reported to be randomized local search heuristics (Brandenburg and Hanauer, 2011; Simpson et al., 2016). This relation with the Feedback Arc Set is important for our method, because it drives the way we provide good solutions by heuristic algorithms.

### Computational complexity

We first check that our problem is also computationally hard. As we have a species tree with infinite weight arcs, we are not in the general case of the Feedback Arc Set problem, so the NP-completeness of our variant is not immediate. However it is easy to reduce the Feedback Arc Set to our problem, leading to the NP-hardness result.

#### Theorem 1

The Maximum Time Consistency problem is NP-hard.

*Proof.* Let us consider an arbitrary instance of the Feedback Arc Set in the form of a weighted graph with *n* vertices (Figure 3 (a)). Construct a species tree with 2*n* leaves, connected by *n* cherry nodes (i.e. nodes having two leaves as children), and complete the rest of the tree by any tree topology (for example a comb, Figure 3 (b)). The cherry nodes are identified with the nodes of the graph, so that any arc can be assimilated to a constraint, and a ranked species tree maximizing the set of consistent constraints yields a total order of the vertices of the initial graph maximizing the consistency with the arcs. Any algorithm finding a maximum time consistent set of constraints, applied on the comb with cherries, would find the solution to the feedback arc set. This proves NP-hardness of the Maximum time Consistency problem.

**Figure 3:**
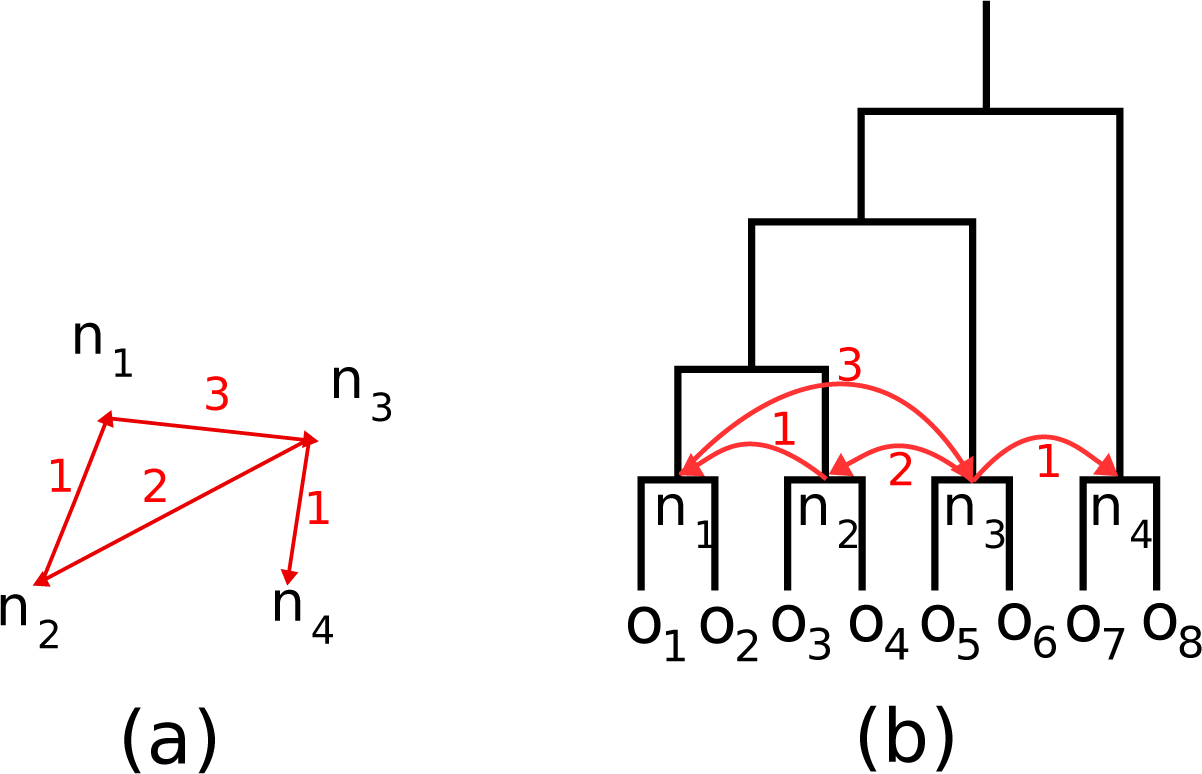
The reduction of Feedback Arc Set to the Maximum Time Consistency problem. (a) an arbitrary instance for the Feedback Arc Set. (b) The instance of Maximum Time Consistency. An algorithm ordering the internal nodes of the species tree will find an optimum feedback arc set.

**Figure 4:**
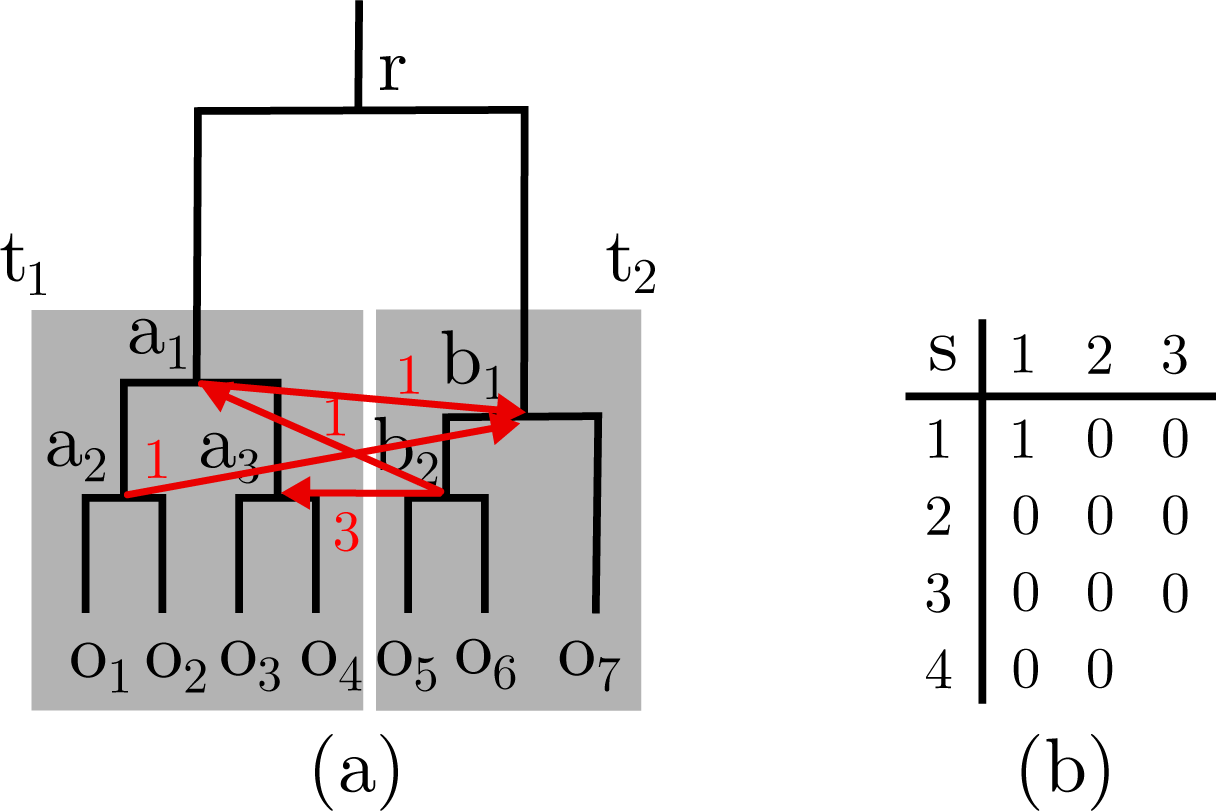
An illustration of the mixing principle. (a) The subtrees a and b are ranked, and there are four constraints between nodes a_1_, a_2_, a_3_ on one side and b_1_, b_2_ on the other. (b) The dynamic programming matrix, with a_1_, a_2_, a_3_ as rows (and an additional row to start the recursion), and b_1_, b_2_ as columns (and an additional column to start the recursion). Since b_2_ has no incoming edge, i.e. incoming(i, b_2_) = 0, s(i, 2) = 0 whatever i. We compute s(3, 1) = s(3, 2) = 0 because b_1_ has no incoming edge from a_3_, then s(2, 1) = s(3, 1) = 0 because a_2_ has no incoming edge. Eventually s(1, 1) = s(2, 1) + incoming(a_1_) = 0 + 1 = 1, so the ordering of the whole tree is r, a_1_, a_2_, b_1_, b_2_, a_3_ and it costs 1 because the constraint b_2_ → a_1_ is not compatible with this order.

### A heuristic principle based on divide-and-conquer approximations

Specific features of our problem, compared to the Feedback Arc Set, do not change the theoretical complexity but can be harnessed to design adapted heuristics. An approximation algorithm to the general Feedback Arc Set problem, which achieves ratio log^2^ *n*, where *n* is the size of the graph, can be obtained by a divide and conquer strategy (Leighton and Rao, 1988). First the graph is cut into two balanced parts. The problem is recursively solved on the two parts and then the two sub-solutions are mixed. The approximation ratio has been improved to *O*(log *n* log log *n*) with similar techniques (Even et al., 1998). The presence of an underlying tree for the graph (the species tree) provides a "natural" way to recursively cut the graph into two. Indeed, let *r* be the root of the species tree (*r* is always the highest node in any ranked tree). Then define *a* and *b* the two subtrees rooted at the two children of *r*. Define three sets of constraints: those having two extremities in *a*, those having two extremities in *b*, and those having one extremity in *a* and one in *b*. The subtree *a* and the first set of constraints, as well as the subtree *b* and the second set of constraints, define new instances of the problem. So the divide step aims at solving independently and recursively the problem on these two instances. This results in ranked trees for *a* and *b*, that is, two independent total orders of the internal nodes of *a* and *b*. Constructing an order of all the internal nodes, that is, containing *r*, the internal nodes of *a* and the internal nodes of *b*, according to the third set of constraints, is the role of the mixing (conquer) step. This is formally described by Algorithm 1.

#### Algorithm 1 Heuristic for Maximum Time Consistency

~~~
1: **procedure** MaxTiC(*r S*, *S*, *C*)
2:   **if** *r* has only leaf descendants **then**
3:        return (*r*)
4:   **else if** *r* has one leaf descendant and one internal node descendant *r*_1_ **then**
5:          Let *C*_1_ be set subset of constraints involving descendants of *r*_1_.
6:          Let (*a*_1_,…, *a*_*k*_) be the result of MaxTiC(*r*_1_, *S*, *C*_1_)
7:          return (*a*_1_,…, *a*_*k*_)
8:   **else**
9:          Let *r*_1_ and *r*_2_ be the children of *r*
10:         Let *C*_i_ be set subset of constraints involving descendants of *r*_*i*_.
11:         Let (*a*_1_,…, *a*_*k*_) be the result of MaxTiC(*r*_1_, *S*, *C*_1_)
12:         Let (*b*_1_,…, *b*_*l*_) be the result of MaxTiC(*r*_2_, *S*, *C*_2_)
13:         Let *C*_inter_ be the subset of constraints involving one descendant of *r*_1_ and one of *r*_2_
14:         Return (*r*)+ the result of MixRank(*S*, *a*_1_,…, *a*_*k*_, *b*_1_,…, *b*_*l*_, *C*_*inter*_)
15:  **end if**
16: **end procedure**
~~~

### The mixing principle

The mixing step of the algorithm consists in obtaining a ranked tree from two ranked subtrees. Note that this procedure can be applied to general approximation algorithms for the Feedback Arc Set. In Leighton and Rao (Leighton and Rao, (1988), the mixing step for the Feedback Arc Set was achieved by simply concatenating the two orders obtained from the solutions to the two subproblems. We propose here a better (optimal) way to achieve this mixing by dynamic programming. Note that our method improves on the approximation solutions to the general Feedback Arc Set problem; however, the approximation ratio however is not improved. Our way to divide the tree into two subtrees even does not guarantee the log^2^ *n* approximation ratio unless the tree and the constraints are balanced enough. We keep this way of dividing despite the lesser theoretical performance because it is the occasion to describe the solution to a more general problem in phylogenetic dating, that we can call Mix ranks, defined as follows. Suppose we are given a rooted binary tree *S*, with root *r*, where the two subtrees *a* and *b* rooted by the children of *r* are ranked. It can be a common situation where two disjoint clades have been dated independently by any method, including, but not limited to, a recursive application of the divide and conquer principle. Suppose also that (possibly conflicting) relative time constraints between internal nodes of *a* and *b* are given, which can be the result of lateral gene transfer detection, but possibly any kind of chronological constraint. Then we have to construct a ranked tree for *S* that contains the input ranks of the children subtrees of the root, and is compatible with a maximum weight subset of time constraints.

The algorithm described below proves that this particular situation of the Feedback Arc Set can be solved in polynomial time.

#### Theorem 2

*Mix ranks* can be solved in time O(n^2^m), where n is the number of nodes of the species tree and m is the number of constraints.

*Proof.* The algorithm follows a dynamic programming principle. Let *a*_1_,…, *a*_*k*_, resp. *b*_1_,…, *b*_*l*_, denote the sequence of internal nodes of *a*, resp. *b*, decreasingly ordered by their position in the ranked subtree (by convention *a*_1_ and *b*_1_ are the oldest nodes). Let *C* denote the set of weighted constraints between internal nodes of *a* and *b*. Given a subset *N* of the internal nodes of the species tree, note *C*_N_ the set of constraints which have both their extremities in *N*.

Let *N*_*ij*_ = {*a*_i_,…, *a*_*k*_, *b*_*j*_,…, *b_l_*}, and let *s*(*i*, *j*) be the minimum sum of the weights in a set which has to be removed from *C*_Nij_, in order to get a consistent set, also compatible with the orders *a*_*i*_,…, *a*_*k*_ and *b*_*j*_,…, *b*_*l*_. It is easy to see that the value of the optimal solution to Mix ranks is *s*(1, 1). We compute it recursively with the following equations,

- *s*(*k* + 1, *j*) = *s*(*i*, *l* + 1) = 0 for all *i ≤ k* and *j ≤ l*,
- *s*(*i*, *j*) = min(*s*(*i* + 1, *j*) + *incoming*(*a*_*i*_, *j*), *s*(*i*, *j* + 1) + *incoming*(*i*, *b*_*j*_)), if *i ≤k* and *j ≤l*,

where *incoming*(*a*_*i*_, *j*) is the total cumulative weight of constraints starting with *b*_*j*_…*b*_*l*_ and ending on *a*_*i*_, and *incoming*(*i*, *b*_*j*_) is the total cumulative weight of constraints starting with *a*_*i*_…*a*_*k*_ and ending on *b*_*j*_.

This translates into a dynamic programming scheme: fill the *s*(*i*, *j*) matrix by using for each cell the adjacent ones. Backtracking along the matrix of *s*(*i*, *j*) gives the optimal mixing of the two orders *a*_1_,…, *a*_*k*_ and *b*_1_,…, *b*_*l*_. Putting *r* before the mixed order gives an optimal solution. A formal description is provided see Algorithm 2.

##### Algorithm 2 Exact algorithm for mixing ranks

~~~
1: **procedure** MixRank(*S*, *a*_1_,…, *a*_*k*_, *b*_1_,…, *b*_*l*_, *C*)
2:  Let *s*(*k* + 1, *j*) = *s*(*i*, *l* + 1) = 0 for all *i ≤ k* and *j ≤ l*
3:  **for** *i* = *k* downto 1, and *j* = *l* downto 1 **do**
4:    Compute *incoming*(*a*_*i*_, *j*) as the sum of all constraints from some *b*_*j*_…*b*_*l*_ to *a*_*i*_
5:    Compute *incoming*(*i*, *b*_*j*_) as the sum of all constraints from some *a*_*i*_…*a*_*k*_ to *b*_*j*_
6:    **if** *s*(*i* + 1, *j*) + *incoming*(*a*_*i*_, *j*) *< s*(*i*, *j* + 1) + *incoming*(*i*, *b*_*j*_) **then**
7:       *s*(*i*, *j*) = *s*(*i* + 1, *j*) + *incoming*(*a*_*i*_, *j*)
8:       *back*(*i*, *j*) = *a*_i_
9:     **else**
10:       *s*(*i*, *j*) = *s*(*i*, *j* + 1) + *incoming*(*i*, *b*_*j*_)
11:       *back*(*i*, *j*) = *b*_j_
12:    **end if**
13:  **end for**
14:  Let *i ←*1, *j ←*1, Result ←∅ *t>* Here begins the backtracking
15:  **while** *i ≤k* and *j ≤l* **do**
16:    Result ←back(*i*, *j*) + Result
17:    **if** *back*(*i*, *j*) = *a*_*i*_ **then**
18:      *i* = *i* + 1
19:    **else**
20:      *j* = *j* + 1
21:  **end if**
22:  **end while**
23:  **if** *i ≤k* **then**
24:    Return Result + (*a*_*i*_,…, *a*_*k*_)
25:  **else**
26:     Return Result + (*b*_*j*_,…, *b*_*l*_)
27:  **end if**
28: **end procedure**
~~~

The time complexity depends on the computation of *incoming*(*a*_*i*_, *j*) which takes at most *O*(*m*) operations, and is called at most *O*(*n*^2^) times, once for all *s*(*i*, *j*). So the running time is bounded by *O*(*n*^2^*m*).

The time complexity depends on the computation of *incoming*(*a*_*i*_, *j*) which takes at most *O*(*m*) operations and it is called at most *O*(*n*^2^) times, once for all *s*(*i*, *j*). So the running time is bounded by *O*(*n*^2^*m*).

Applying the mixing algorithm as a conquer step yields a recursive heuristic for the general problem, which consists in applying Algorithm 1 with parameters *r*, *S*, *C*, where *r* is the root of the species tree *S* and *C* is the set of time constraints.

### Implementation

In our software MaxTiC, we implemented in Python the heuristic recursive principle described above. In practice the running time is almost instantaneous for all the simulated datasets we tested. We also implemented two other heuristics: a greedy heuristic and a local search approach.

The greedy heuristic consists in sorting the constraints in decreasing order of their weight, and examine them one by one in that order. Each constraint is kept in a consistent set if it is compatible with the partial order given by the species tree and not conflicting with other constraints already marked as kept, and discarded otherwise.

The local search consists in performing a randomized hill-climbing in the space of linear extensions of the partial order given by the species tree, that is, total orders on internal nodes that do not contradict the partial order. From one of these total orders, the algorithm chooses one element (internal node) uniformly at random, and changes its position to an alternative one, chosen uniformly at random among all possible positions. The obtained total order is the proposition. The algorithm accepts it as the new state if it is compatible with the partial order given by the species tree and if it does not decrease the cumulative weight of the compatible constraints, compared with the current state. The search is run for a prescribed time given as a parameter by the user, and this is its only way to terminate.

We tested this program on simulated data, taking the best solution out of the greedy one and the heuristic one, and applying on it the local search for a fixed run-time of three minutes.

## IV. Results

### Transfer rate and number of inferred transfers

We first tested the ability of the ALE method to infer a likely number of transfers, as well as the effect of inferring transfers in a phylogeny which is a small subtree of the one on which transfers have been simulated. On Figure 5, we can see that up to a very high transfer rate, the number of inferred transfers follows a regular function of the transfer rate. Measures of transfer numbers on biological datasets were done for comparison purposes from the cyanobacteria and fungi dataset from Szöllősi et al. (2015). They show that the range of the simulation parameters contains numbers of transfers per family comparable to published biological datasets.

**Figure 5:**
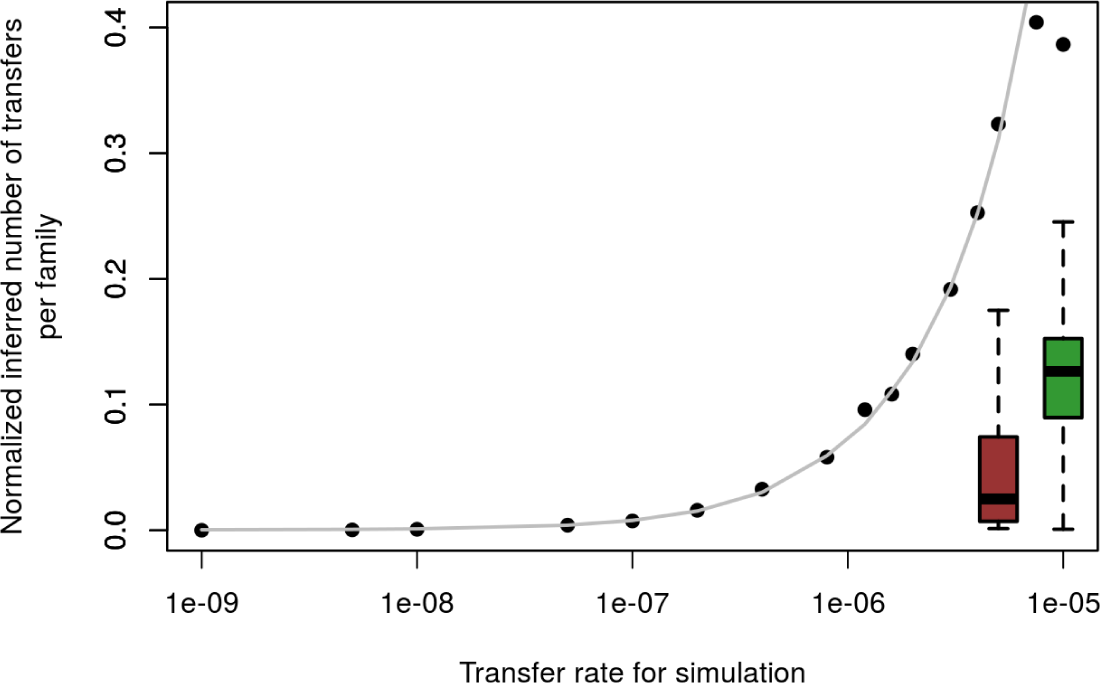
Mean number of inferred transfers (number of transfers per family and per branch of the species tree), as a function of the transfer rate in the simulation (*log*_10_ scale). Each point is one simulation of a species tree and 1000 gene trees with its own transfer rate. The right boxplots show the distribution of the number of inferred transfers on gene families from two published biological datasets: 28 Fungi (red) and 40 cyanobacteria (green) (Szöllősi et al., 2015). For each gene family the number of inferred transfers per branch is computed. It shows that comparable numbers are found in simulated and biological datasets.

Henceforth, we use the mean number of inferred transfers as the reference instead of the transfer rate, to relate our measures to numbers comparable with what is found in biological datasets, for which we do not know the transfer rate on the complete phylogeny containing extinct and unsampled species.

### Number of conflicting constraints

We then measured the fraction of constraints that has to be discarded to get a consistent set of constraints (Figure 6). We compare this value to the fraction of the constraints not compatible with the true (simulated) ranked species tree (red points). We see that the values on reconstructed node orders are close and always a bit under the true values. This justifies the minimizing approach: the true conflict is close to the minimum. However as the optimum is always lower than the true value, it also shows that discrepancies are not due to limitations in the optimization algorithm but to limitations in the model itself. Another lesson to be drawn from this figure is that for what seem to be biologically relevant transfer rates, between 5% and 20% of constraints must be removed to get a consistent subset. This means that at least this amount of transfer is wrongly inferred and this places a lower bound on the rate of false positive transfers output by ALE. It has already been observed that current transfer detection methods usually infer an accurate number of transfers but they are less precise for the identification of donors and recipient (Abby et al., 2010).

**Figure 6:**
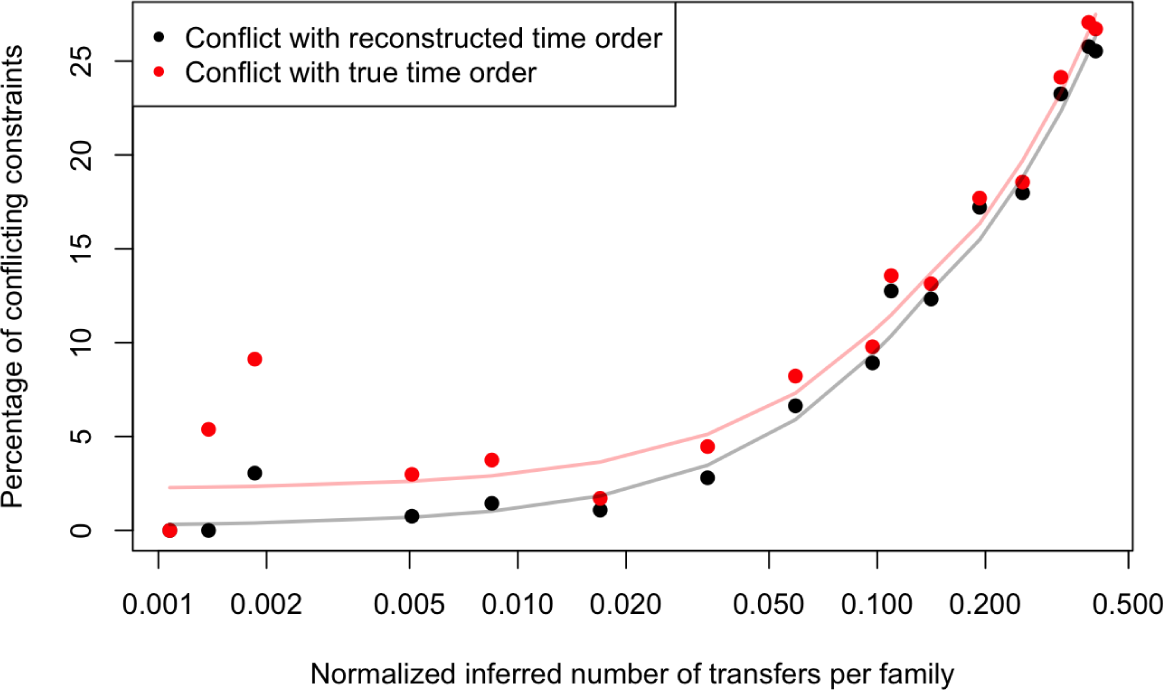
Fraction of constraints that have to be removed in order to get a consistent set, as a function of the mean number of inferred transfers (*log*_10_ scale). Red dots are for the fraction of constraints in conflict with the true (simulated) tree, and black dots are for the fraction of constraints in conflict with the reconstructed tree, minimizing the conflicts. Horizontal boxplots show the number of inferred transfers from two biological datasets: 28 Fungi (red) and 40 cyanobacteria (green) (Szöllősi et al., 2015).

### Similarity between inferred and true ranked trees as a function of the number of gene families

We give an idea of how many gene trees (and as a consequence how many transfers) are necessary to obtain an accurate ranking information. In Figure 7 (bottom), we plot the Kendall similarity between the true tree and the obtained tree, as a function of the number of gene trees, for a constant transfer rate of 1.6 × 10^−6^, corresponding to approximately 5 inferred transfers per family (all families have approximately 100 genes).

**Figure 7:**
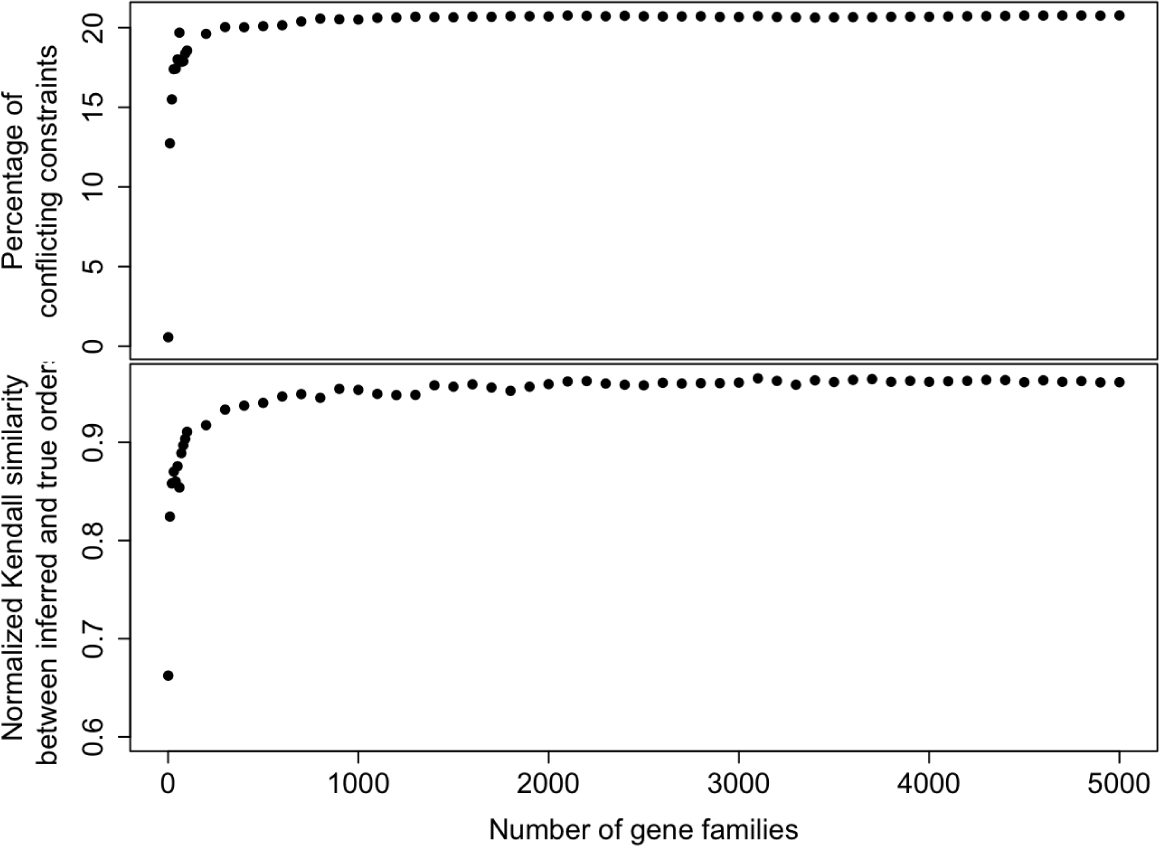
Top: Fraction of the constraints removed by MaxTiC to get a consistent set as a function of the number of gene trees. Bottom: Normalized Kendall similarity of the true ranked tree and the obtained ranked tree, as a function of the number of gene trees in the experiment.

We can see that the method starts with a very low similarity if there are not enough gene trees, which is expected as in the absence of transfers there is no information to infer the ranked tree. Then the similarity rapidly increases, almost reaching a plateau at about 400 families, then slowly increasing up to 5000. This means that the more gene trees are available, the better the result will be, but with only marginal gains in accuracy after 1000 gene trees. On the top panel of the Figure 7, we see that the conflict (ratio of removed constraints to obtain a consistent set) also grows quickly and then stays remarkably stable. This shows that the rate of signal and conflict is relatively constant in all families.

### Sensitivity to the transfer rate

We then investigated the effect of the transfer rate on the accuracy of the ranking. We measured the normalized Kendall similarity as a function of the average number of transfers per gene family. The results are shown on Figure 8. As expected, too few transfers result in a somewhat inaccurate result, because of a lack of signal, and too many transfers make the similarity to the true ranking decrease. However the slopes are very different: whereas a reasonable number of transfers are sufficient to give a good ranked tree, the ranked tree stays reasonably good even with a huge number of transfers (several dozens per family).

**Figure 8:**
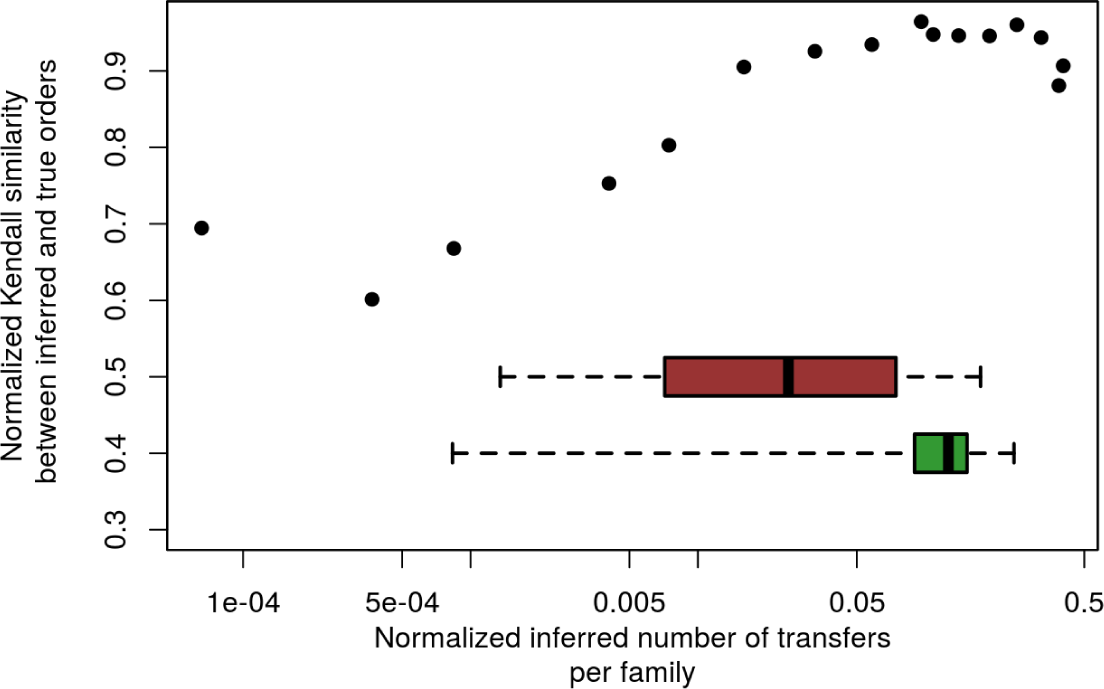
Normalized Kendall similarity of the true ranked tree and the obtained ranked tree, as a function of number of transfers, per branch and per family (*log*_10_ scale). The boxplots show the distribution of the number of inferred transfers on gene families from two published biological datasets: 28 Fungi (red) and 40 cyanobacteria (green) (Szöllősi et al., 2015). For each gene family the number of inferred transfers per branch is computed.

Note, however, that in all cases considered, the normalized Kendall similarity to the true ranked tree remains bounded from above at around 95%, and under almost all conditions, in particular conditions that produce numbers of transfers consistent with those observed in Fungi and Cyanobacteria, it is between 90% and 95%. This implies that it is possible to use ALE_undated to detect transfers and get a result close to the real order of speciations in a wide range of conditions.

Note, finally, that the amount of conflict (that can be measured on real data) is not necessarily a good proxy for the similarity (that requires the knowledge of the true ranked tree): as shown by a comparison of Figures 6 and 8, the behaviours of the two variables have no evident correlation when the transfer rate increases.

### Sensitivity to non modeled processes and errors in the gene trees

We examine the effect of non modeled processes and gene tree errors (Figure 9). In SimPhy it is possible to vary the population size, and with the population size the probability of incomplete lineage sorting (ILS) increases. ALE does not model ILS, thus any deviation from the species tree topology resulting from ILS will be interpreted as a series of DTL events. Indeed, it can be seen on Figure 9 (middle) that for a same transfer rate, the number of inferred transfers increases with population size, thus with the amount of ILS. On The top panel of Figure 9 shows that these spurious transfers are not time compatible as the frequency of conflicting transfers increases also with population size. However on the bottom panel, we can see that nonetheless, and despite a decrease in the Kendall similarity with the true ranked tree, it is still possible to obtain a reasonable ranking of the species tree even in the presence of a high rate of false transfers that results from ILS or phylogenetic error.

**Figure 9:**
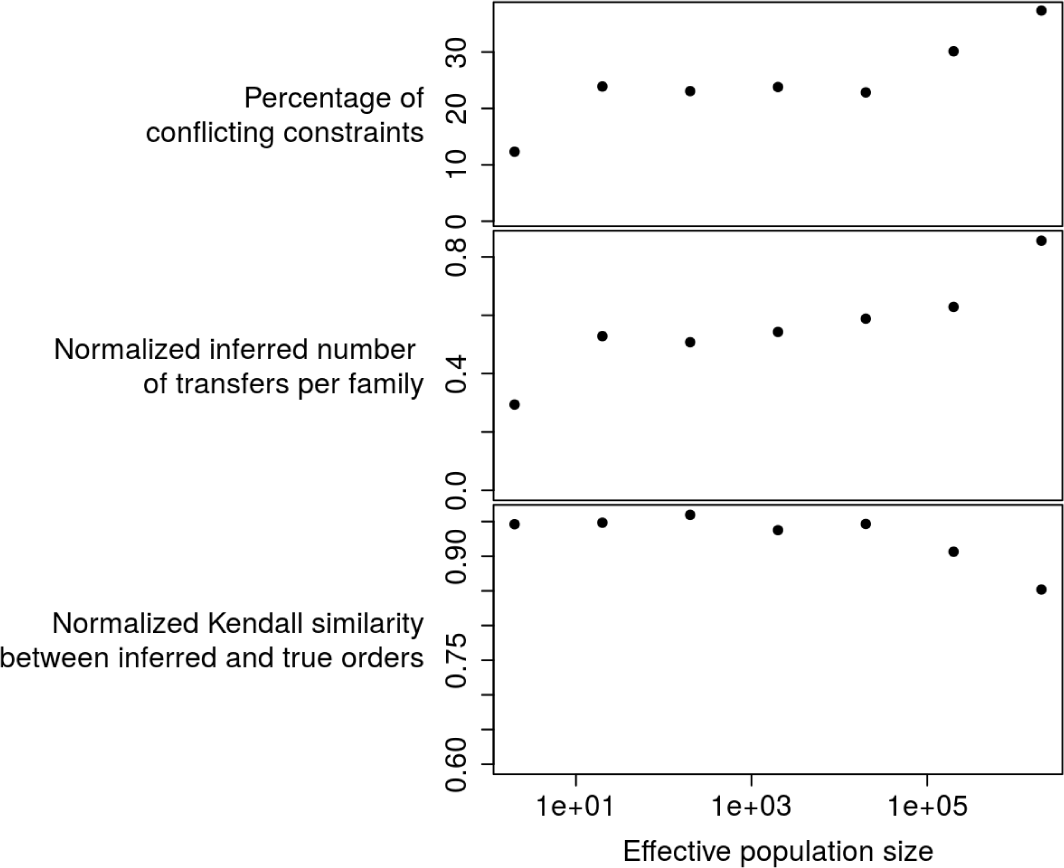
Minimum fraction of conflicting constraints, mean number of inferred transfers per family and normalized Kendall similarity of the true ranked tree and the obtained ranked tree, as three functions of population size (*log*_10_ scale), for a fixed transfer rate (10^−6^ in the simulation). Population size favors incomplete lineage sorting in SimPhy; as such, it is used here to measure the effect of non modeled processes or as a proxy for errors in phylogenetic reconstruction.

### Sensitivity to errors in the species tree

Finally, the topology of the species tree is in general not known with high precision, so we tested the robustness of the method to errors in the species tree. We chose a simulated dataset with a transfer rate of 10^−6^, because this is one of the rates that lead to a number of transfers per family close to what we measured on the two biological datasets. Then we compared the normalized Kendall similarity of 5 simulations with the true species tree (blue dots in Figure 10), with 5 simulations for 5 different conditions: re-rooting the species tree at a grand-child of the root (red dots), and respectively applying 5, 10, 15 and 20 random “nearest neighbor interchanges” (NNIs) to the species tree (green dots). We plot in Figure 10 the normalized Kendall similarity in function of the obtained Robinson-Foulds distance to the true tree (A certain number of random NNIs leads to a Robinson-Foulds distance of at most this number).

**Figure 10:**
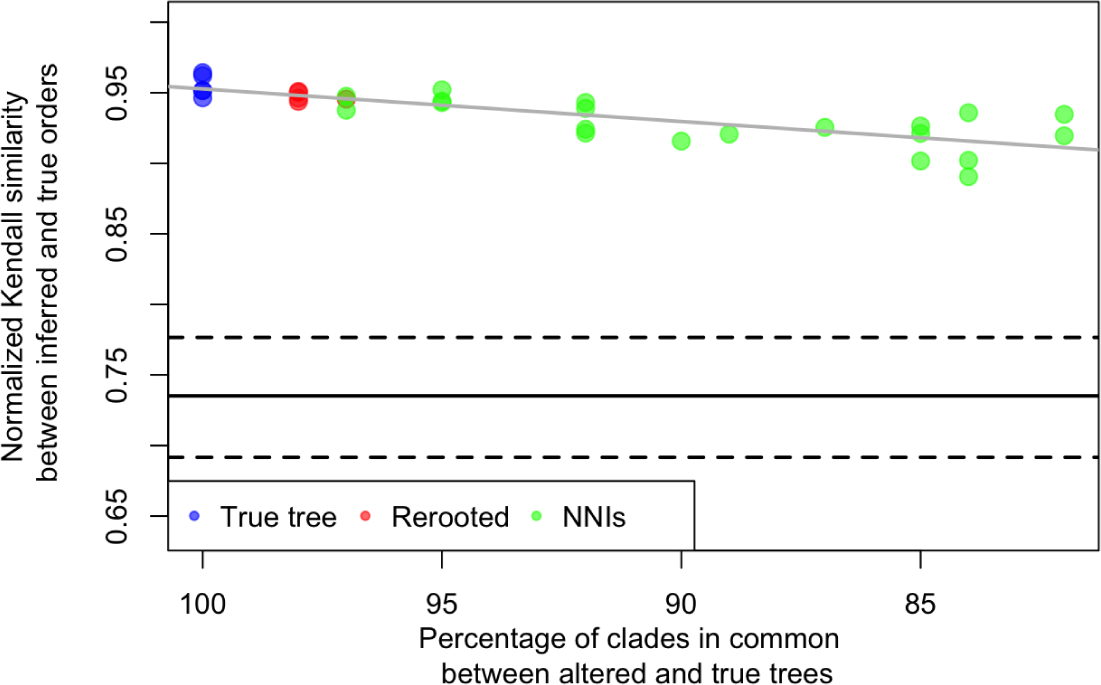
Normalized Kendall similarity of the true ranked tree and the obtained ranked tree, for different species trees in which errors have been introduced. The Kendall similarity is computed on the common clades, that is, on the fraction of true clades in the modified species tree. The three horizontal lines show three quantiles of the distribution of normalized Kendall similarity of randomly generated ranked trees. This shows that transfers give a robust dating information even in the presence of a highly erroneous species tree.

The tendency of Figure 10 shows a good robustness to errors in the species tree, showing that even with quite distant species trees, the rank of true clades is well preserved.

### Application to a small Cyanobacteria dataset

In order to test the ability of the program to find good solutions in biological conditions, and to test the dating method on a tree coming from biological data, we examined the Cyanobacteria transfers presented in Szöllősi et al. (2015) on a subtree restricted to 14 leaves out of the 40 in the original study. This restriction has the purpose to be able to enumerate all possible rankings (there are 69300 of them on this tree) and to test our heuristic against an exhaustive solution search. This example is described into details, with all associated data, in the maxtic repository https://github.com/ssolo/ALE/tree/master/maxtic.

We found that 35% of the total constraint weight has to be discarded to get a consistent set of time constraints. This is similar to the worst conditions in our simulations, with the largest contribution of non-modeled processes in the gene trees. This is not surprising since it is known that with real sequences phylogenetic errors are not rare. This number corresponds to both the optimal value that was found using the exhaustive search, and the value given by the Maximum Consistency Heuristic, so it is a good sign that we can find decent solutions in instances coming from a biological dataset.

A study of the full 40 leaf tree, along with comparisons with other domains of life, tests of the transfer dating methods with molecular clocks, can be found in a companion paper (Davin et al., 2017).

## V. Conclusion

In this paper, we give a proof of principle of a method to get a ranked species tree with the information provided by gene transfers. We present a method and a software, called MaxTiC for Maximum Time Consistency, taking an unranked species tree as input, together with a set of possibly conflicting weighted time constraints, and outputting a ranked tree maximizing the total weight of a compatible subset of constraints. We validate our methodology on simulations produced by an independent software SimPhy, with characteristics that we compare to published biological datasets. The results confirm that we can date with transfers under a wide range of conditions including errors in gene trees and species trees. This additional source of information for dating can be a good alternative to fossils and the (relaxed) molecular clock since the fossil record is poor or difficult to interpret precisely in clades where transfers are abundant.

The scores of the solutions are informative about the inference of trees, transfers and dates. The 5% to 35% conflict in constraint sets tells us that there is at least this amount of false positive at the level of time constraints. In biological data we can invoke clustering sequences into families, and gene tree reconstructions to explain part of the error. However here on simulated data we control for these and the false positive rate, while lower, remains important. This could also be explained by uncertainties or errors in reconciliation scenarios. A more conservative way of transforming transfers into time constraints, which would give less weight to particular reconciliations can be proposed: if, back to Figure 1, we detect a transfer between *A* and *B* but *X* or *Y* are not represented in the associated gene tree, the definition of the time constraint could be relaxed to the first represented ancestor of *A*, and first represented descendants of *B*. In our experience, such coding approximately yielded half as much false positives, but without improving time order inference.

This points to an interesting byproduct of our method and analyses. MaxTiC is able to filter out a set of transfers detected by phylogenetic methods and detect false positive. Even if the false positive rate is high, MaxTiC produces good rankings, meaning that besides dating, it can be used to discriminate *bona fide* transfers from artefactual ones.

## VI. Acknowledgments

G.J.Sz. received funding from the European Research Council (ERC) under the European Union’s Horizon 2020 research and innovation programme under grant agreement No. 714774. This project was supported by the French Agence Nationale de la Recherche (ANR) through grant no. ANR-10-BINF-01–01 ‘Ancestrome’.

